# XplainTDA: Explaining the topological structure of single-cell multi-omics data through perturbation

**DOI:** 10.64898/2026.10.14.759129

**Authors:** Isabel Giray, Gabriele Malagoli, Katharina T. Schmid, Maria Colomé-Tatché

**Affiliations:** Department of Mathematics, TUM School of Computation, Information and Technology, Technical University of Munich, Munich, Germany; Biomedical Center, Division of Physiological Chemistry, Faculty of Medicine, Ludwig-Maximilians-Universität Munich, Munich, Germany; Institute of Computational Biology, Computational Health Center, Helmholtz Munich, Munich, Germany; Hospital del Mar Research Institute (HMRIB), Barcelona, Spain

**Author notes:** These authors contributed equally to this work (Isabel Giray, Gabriele Malagoli).

## Abstract

Single-cell experiments have improved our ability to study biological systems by profiling thousands of cells simultaneously. The data can be viewed as a point cloud in a proper experimental space, whose shape is defined by the gene regulatory network. Persistent homology has been proposed as a method for studying the intrinsic structure of data and has shown the potential to reveal novel insights. Existing approaches focus on detecting topological features or groups of cells, without a focus on explainability and interpretability. Here, we introduce an explainable topological data analysis framework for single-cell data (XplainTDA) that identifies molecular determinants underlying the topological structure of the data via a perturbation-based approach. Across omics and data sets, we show that the method is stable, reproducible, and interpretable, extending persistent homology from topological description to molecular interpretation.

## 2 Background

Single-cell sequencing has transformed molecular biology by enabling high-throughput characterisation of genomic, transcriptomic, and epigenomic states at single-cell resolution. The increasing availability of robust and standardised experimental workflows has accelerated the adoption of these technologies across a wide range of biological applications. For example, single-cell RNA sequencing (scRNA-seq) [1], which profiles gene expression (GEX), and single-cell Assay for Transposase-Accessible Chromatin using sequencing (scATAC-seq) [2], which measures chromatin accessibility, can be performed using commercially available platforms such as those developed by 10x Genomics. More recently, advances in single-cell multi-omics have enabled simultaneous interrogation of multiple molecular layers within a single cell, including gene expression and chromatin accessibility [3–5]. However, the data are sparse, noisy, and high-dimensional, requiring robust computational methods to extract meaningful biological insights. The first challenge is dimensionality reduction [6]. The ambient space, which is the space defined by the experiment, i.e. the complete set of genes for GEX or peaks for ATAC, has tens or hundreds of thousands of dimensions, which makes it impossible to sample it properly due to the curse of dimensionality [7]. The common assumption in data analysis, known as the manifold hypothesis [8, 9], states that the data actually lie on a lower-dimensional manifold with fewer degrees of freedom than the observed space. Dimensionality reduction methods, such as principal component analysis (PCA) [10], autoencoders (AEs) [11–13], and other nonlinear dimensionality-reduction methods (such as PHATE [14], UMAP [15], and t-SNE [16]), aim to learn a representation of the data in a smaller space. The second challenge with the data is identifying the key degrees of freedom of the system, i.e., the genes or peaks that shape the manifold and define the observed phenotypes (such as cell type, disease state, or other cell annotations). In the single-cell field, the common practice for accomplishing such a task is differential analysis (DA) [6]. Given two groups of cells, such as two different cell types, DA identifies the features that follow a statistically different distribution between them. Other approaches rely on topic modelling (TM) [17, 18], grouping features by co-expression in cells, or explainable AI (XAI) [19], which extracts the most useful features for classification tasks based on their impact on predictions.

In recent years, topological data analysis (TDA), specifically persistent homology (PH) [20–23], has been shown to be a useful approach for investigating biomedical data, from single-cell RNA-seq to functional MRI (fMRI) [24–31]. PH characterises the topological structure of data across multiple spatial scales and summarises the appearance and disappearance of topological features in a persistence diagram (PD). Existing applications of PH to biological data have primarily used these topological summaries to characterise data structure or group observations, without directly relating the resulting topology back to the molecular input features. Here, we introduce a perturbation-based approach to identify the genes and peaks that have the largest influence on the topological structure of single-cell data. By linking changes in persistent topology to individual molecular features, our approach provides a complementary perspective for the interpretation of single-cell data.

## 3 Results

### 3.1 The XplainTDA model

XplainTDA first learns a low-dimensional representation of the single-cell data on which persistent homology is computed. Given an initial count matrix *X* ∈ ℝ^*N×P*^, where *N* is the number of cells and *P* the number of genes or peaks, we embed the data using a geometry regularised autoencoder (GRAE) [9], yielding

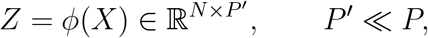

where *ϕ* denotes a GRAE (Fig. 1 A). In previous work, we showed that GRAE provides suitable embeddings of scATAC-seq and scRNA-seq data, particularly in the context of interpretability [19]. Persistent homology is then used to characterise the topology of the embedded point cloud

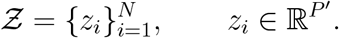

**Fig. 1:**
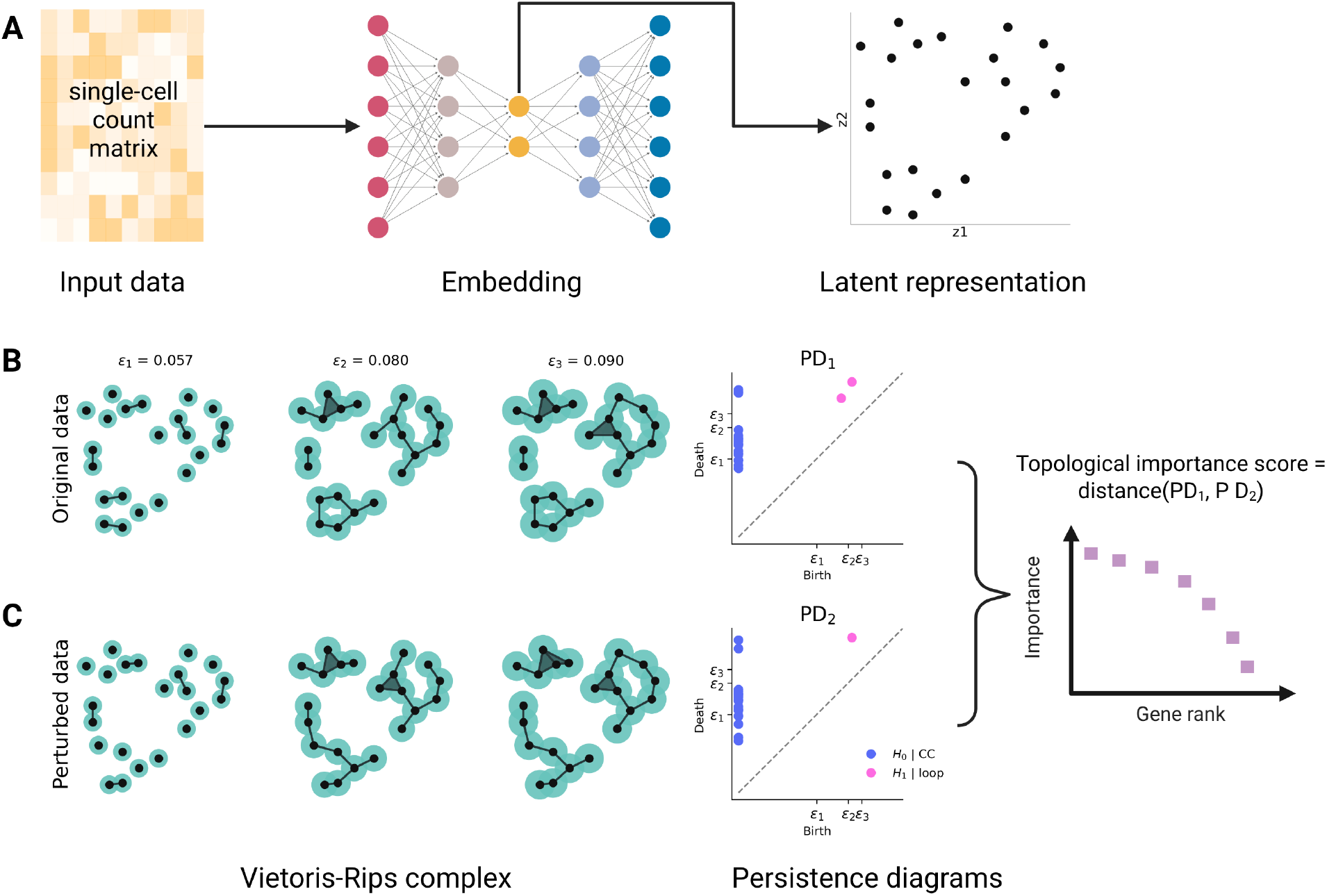
**A** The single-cell data are stored as a count matrix, and their dimensionality is reduced using the geometry regularised AE. **B** Schematic of the Vietoris-Rips complex at different scales and its subsequent persistence diagram. **C** In the perturbed cell representation the model computes PH and finally quantifies the difference between the original and new persistence diagram to assign a topological importance score to each gene (or peak in scATAC-seq data). Figure partially created with BioRender.

For a scale parameter *ϵ* ≥ 0, the Vietoris–Rips complex *R*_*ϵ*_(Ƶ) [32] contains a simplex for every subset of points whose pairwise distances are smaller than or equal to *ϵ*. At small values of *ϵ*, the complex consists mainly of isolated points, whereas increasing *ϵ* progressively introduces edges and higher-dimensional simplices. Vietoris–Rips complexes satisfy the nesting relation

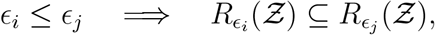

which defines a filtration [22, 23, 33].

At each scale, the homology groups *H*_*h*_ encode topological features of dimension *h*: *H*_0_ represents connected components, *H*_1_ loops or tunnels, and *H*_2_ enclosed voids. As *ϵ* increases, individual topological features can appear and disappear. PH records the scale at which each feature *l* appears, its birth *b*_*l*_, and the scale at which it disappears, its death *d*_*l*_. For each homological dimension *h*, these values form the persistence diagram

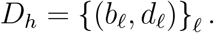

The persistence *d*_*l*_ − *b*_*l*_ measures the range of scales over which a topological feature exists, with longer-lived features persisting over a broader range of the filtration.

Computing the full Vietoris–Rips complex becomes prohibitively expensive for large single-cell data sets. Therefore, XplainTDA constructs a *k*-nearest-neighbour (*k*-NN) graph on *Z* and generates the filtered simplicial complex from this sparse representation. Persistent homology is then computed with *Ripser* [34], yielding a persistence diagram *D*_*h*_ for each homological dimension *h* (Fig. 1 B).

To assess the contribution of individual biological features, either genes or peaks, the input data are perturbed in silico: persistent homology is recomputed after masking a feature *p*, yielding a perturbed persistence diagram (PD) 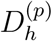. The resulting topological change is quantified by the difference between the original and perturbed persistence diagrams, which defines the feature importance

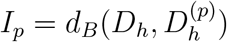

where *d*_*B*_(·, ·) denotes the *q-bottleneck loss* (Fig. 1 C), a variant of the standard bottleneck distance [35]. The quantile makes the loss more robust to noise, ensuring a conservative approach to feature interpretability: genes or peaks that affect only a small part of the topology will be ranked lower. The diameter normalisation helps interpret the values; its simple geometric meaning indicates the topological discrepancy with respect to the spatial scale, namely, the diameter. Features that yield larger values are interpreted as exerting a stronger influence on the topological structure.

For all tests and results, we used three single-cell multi-omic (GEX and ATAC) data sets from three different tissues: peripheral mononuclear blood cells (PBMC) [36], human adult brain cortex (Brain) [37], and bone marrow mononuclear blood cells (BMMC) [38].

### 3.2 Topological structure is robust to k-NN graph construction and cell subsampling

The computational complexity of PH depends strongly on the number of simplices required to construct the Vietoris–Rips complex, which can become computationally infeasible for large data sets such as single-cell data; hence, we approximate it using a *k*-nearest-neighbour (*k*-NN) graph constructed on *Z*. Let *N* be the number of cells and *k* the number of nearest neighbours. For a full Vietoris–Rips complex, the number of *j*-simplices, denoted by 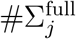, scales as

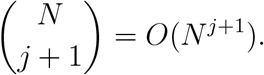

To compute persistent homology in dimension *H*_*h*_, simplices up to dimension *h* + 1 are required, giving 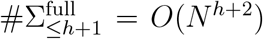. For a *k*-NN sparse clique complex, the number of *j*-simplices scales approximately as 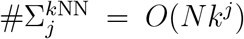, and therefore 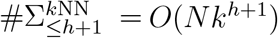. Thus, for homology dimension *H*_*h*_, the scaling changes from *O*(*N*^*h*+2^) to *O*(*Nk*^*h*+1^).

We assessed the effect of this approximation by varying *k* in the *k*-NN graph and comparing the resulting persistence diagram with a reference diagram using the *q*-bottleneck loss. For *H*_0_, we used the persistence diagram obtained from the full-distance (FD) matrix as the reference. For *H*_1_ and *H*_2_, computing the full-distance persistence diagrams for the complete data sets was infeasible even with 200 GB of RAM on our high-performance computing cluster. We therefore used the diagrams obtained with *k* = 1250, the largest value considered, as the reference for these homology dimensions (Methods).

For all three homology dimensions, we observed a value of *k* above which increasing the neighbourhood size no longer substantially changed the reconstructed persistence diagram relative to the corresponding reference (Fig. 2 A–C). *H*_0_ was stable even for very low values of *k*, indicating that the connectivity structure was only weakly affected by the number of neighbours (Fig. 2 A). *H*_1_ and *H*_2_ stabilised at approximately *k* = 15 and *k* = 50, respectively (Fig. 2 B, C). To be conservative, we therefore set *k* = 75 as the default for the subsequent analyses.

**Fig. 2:**
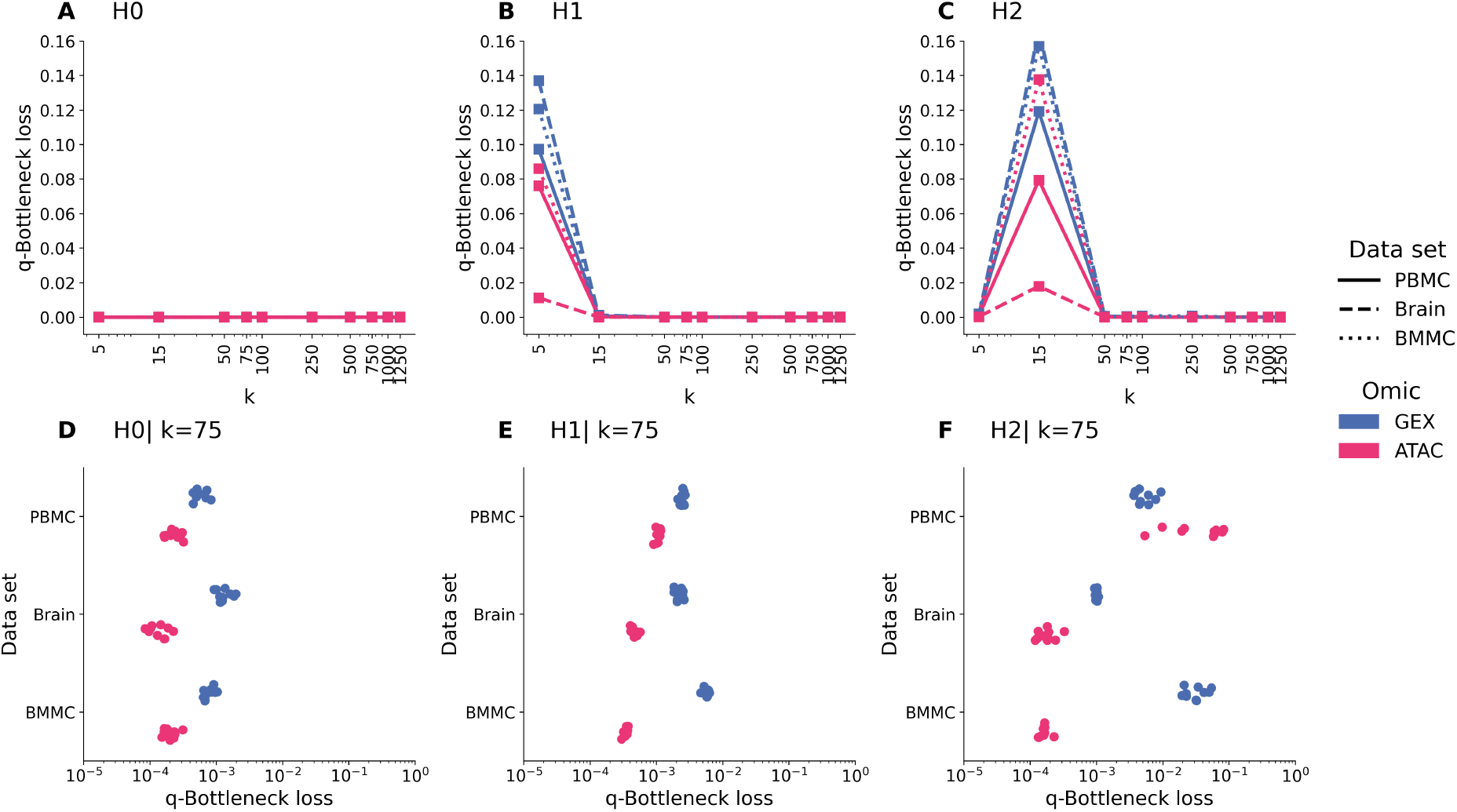
Topological stability of the inferred representations across neighbourhood size and cell subsampling. **A–C** Quantile-normalised *q*-bottleneck loss for homology dimensions *H*_0_, *H*_1_, and *H*_2_ as a function of the number of nearest neighbours *k*. For *H*_0_, persistence diagrams are compared with the diagram obtained from the full-distance graph, whereas for *H*_1_ and *H*_2_ they are compared with the diagram obtained at the largest value considered, *k* = 1250. Line style denotes the data set and colour denotes the omic. **D–F** Distribution of the quantile-normalised *q*-bottleneck loss under repeated cell subsampling for *H*_0_, *H*_1_, and *H*_2_. Each coloured dot represents the *q*-bottleneck loss between two independently generated stratified subsamples, each retaining 25% of the cells from each cell type. Results are grouped by data set and omic. Lower values indicate greater topological stability.

As mentioned in the introduction, the data manifold is formally unknown and it is approximated by a discrete sampling of cells on it. To test whether PH captures the same topological structure across subsets of cells within the same point cloud, for each data set and each omic, we subsampled five groups of cells without replacement and computed PH on each subsample. Finally, we measured the pairwise differences between their persistence diagrams using the normalised q-bottleneck loss (Methods). The similarity across sub-sampling follows a similar stability trend and the *k* − *NN* approximation: the lower the rank of the homology group, the higher the stability of the persistence diagrams. (Fig. 2). In particular, *H*_2_ show a higher dispersion of the similarity than *H*_0_ and *H*_1_ (Fig. 2 D-F). However, all *H*_*h*_, data sets and omics show a consistent low difference, with respect to the embedding diameter, due to sub-sampling (Fig. 2 D-F).

### 3.3 Connected components capture cell-type structure

In conventional single-cell analysis, cell types are typically identified by applying a community detection algorithm such as Leiden [39] to a cell–cell graph, yielding clusters that are subsequently annotated. The *H*_0_ structure of a Vietoris–Rips filtration provides a closely related notion of partitioning, where cells are grouped according to connected-component membership at a given filtration value *ε*. We inspected the persistence diagrams of the three GEX data sets (Fig. 3 A–C), focusing on *H*_0_ to assess whether there exists a filtration scale at which the topology reflects the known cell-type structure. For this purpose, we examined the evolution of the connected components over *ε* in the gene-expression space, which is better suited to comparison with cell-type annotations because these annotations are defined by gene-expression profiles and marker genes. This analysis is intended as a validation of the biological structure captured by *H*_0_, rather than as a method for de novo cell-type discovery.

**Fig. 3:**
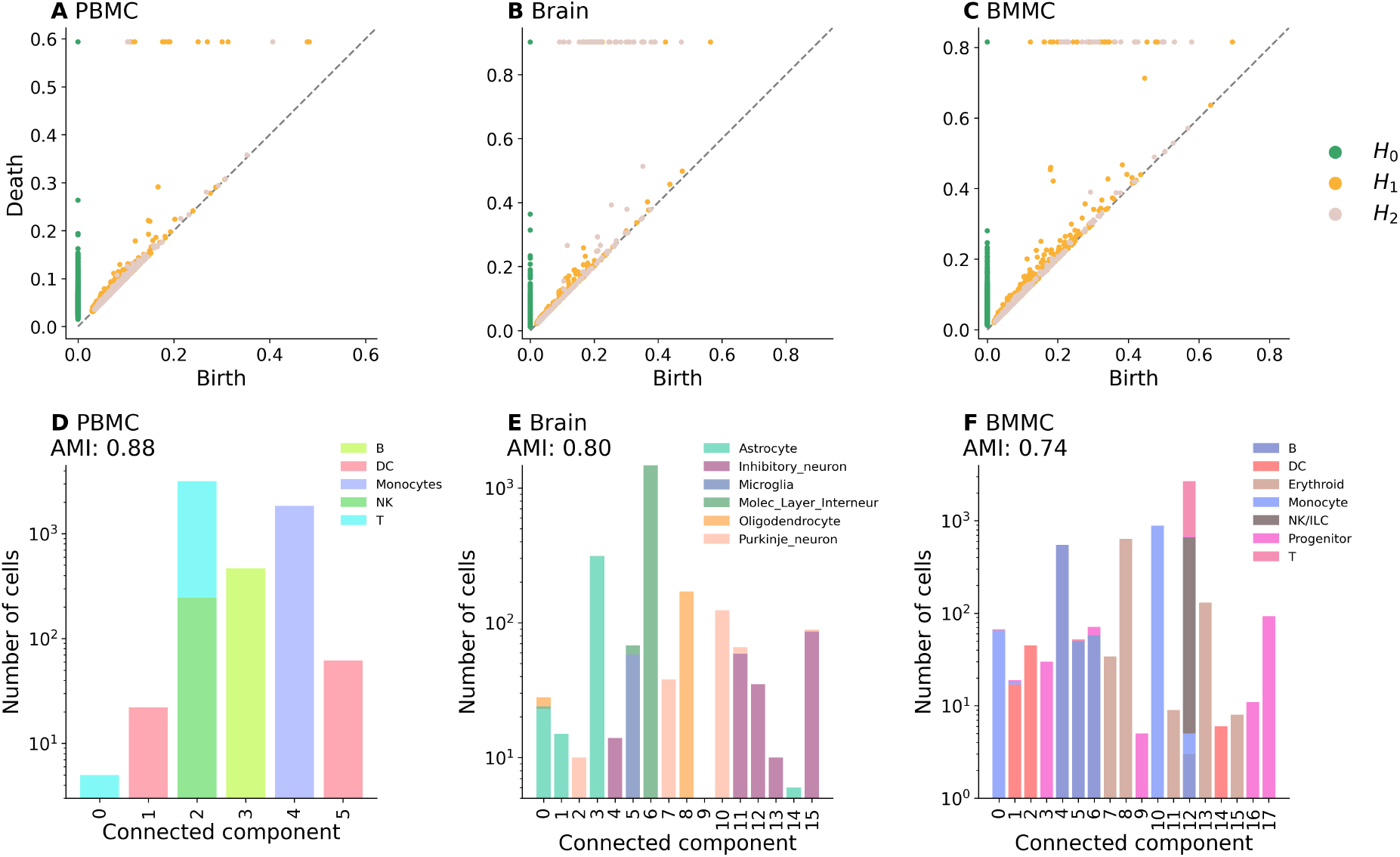
**A-C** Persistence diagrams of the GEX point clouds of the three data sets. **D-F** Correspondence between *H*_0_ topology and reference cell-type annotations at the filtration value showing the maximum AMI. The filtration value is selected using the reference annotations for validation purposes and is not intended as a procedure for de novo cell-type identification.

For all three data sets, a filtration value that resembles the cell-type structure clearly exists (Fig. 3 D, E, F). Visually, all the cell types are correctly separated except for the T and NK cells, which play a similar role in the immune system (Fig. 3 D, F). Quantitatively, the agreement between ground-truth annotations and connected components is evaluated using the adjusted mutual information (AMI) [40]: across all data sets, AMI is consistently high, from 0.74 to 0.88 across the three data sets (Fig. 3 D, E, F).

### 3.4 Perturbation approach to TDA reveals consistent biological marker genes and chromatin motif

We used in silico molecular perturbations to quantify how individual genes and peaks affect the topology of the cell manifold. Let *Z* denote the embedding of the cells, and let *Z*^(*p*)^ denote the embedding obtained after perturbing molecular feature *p*. We write *D*_*h*_(*Z*) for the persistence diagram of an embedding *Z* in homology dimension *h*, and *d* for the normalised quantile bottleneck loss between two PDs. In dimension *h* and for a gene or peaks *p*, the topological importance score is defined as

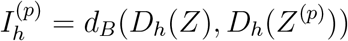

It measures how much the perturbation of the biological features *p* changes the topology of the entire space *Z*, under the lens of the *h* − *th* homology group. This strategy is applied stratifying the cells by annotation, in order to retrieve the importance score of the features for each cell type.

To evaluate the impact on the GEX topology, we select a set of marker genes for each data set and compare the distribution of *I*^(*p*)^ for theese markers with that of a random set of genes of the same size, populated with genes with the same coverage and expression. (Methods). We expect marker genes to have a greater impact on the topology than a random set of genes. Across all homology groups and data sets, marker-gene distributions are indeed shifted towards higher importance scores than the corresponding random controls (Fig. 4). Notably, this pattern was observed not only for *H*_0_ (Fig. 4 A, D, G), which reflects the connectivity of the *k*-NN graph and partitions cells into connected components, but also for the higher homological dimensions *H*_1_ and *H*_2_ (Fig. 4 B, C, E, F, H, I). This indicates that higher-order topological structure retains meaningful information about biological features relevant to cellular identity.

**Fig. 4:**
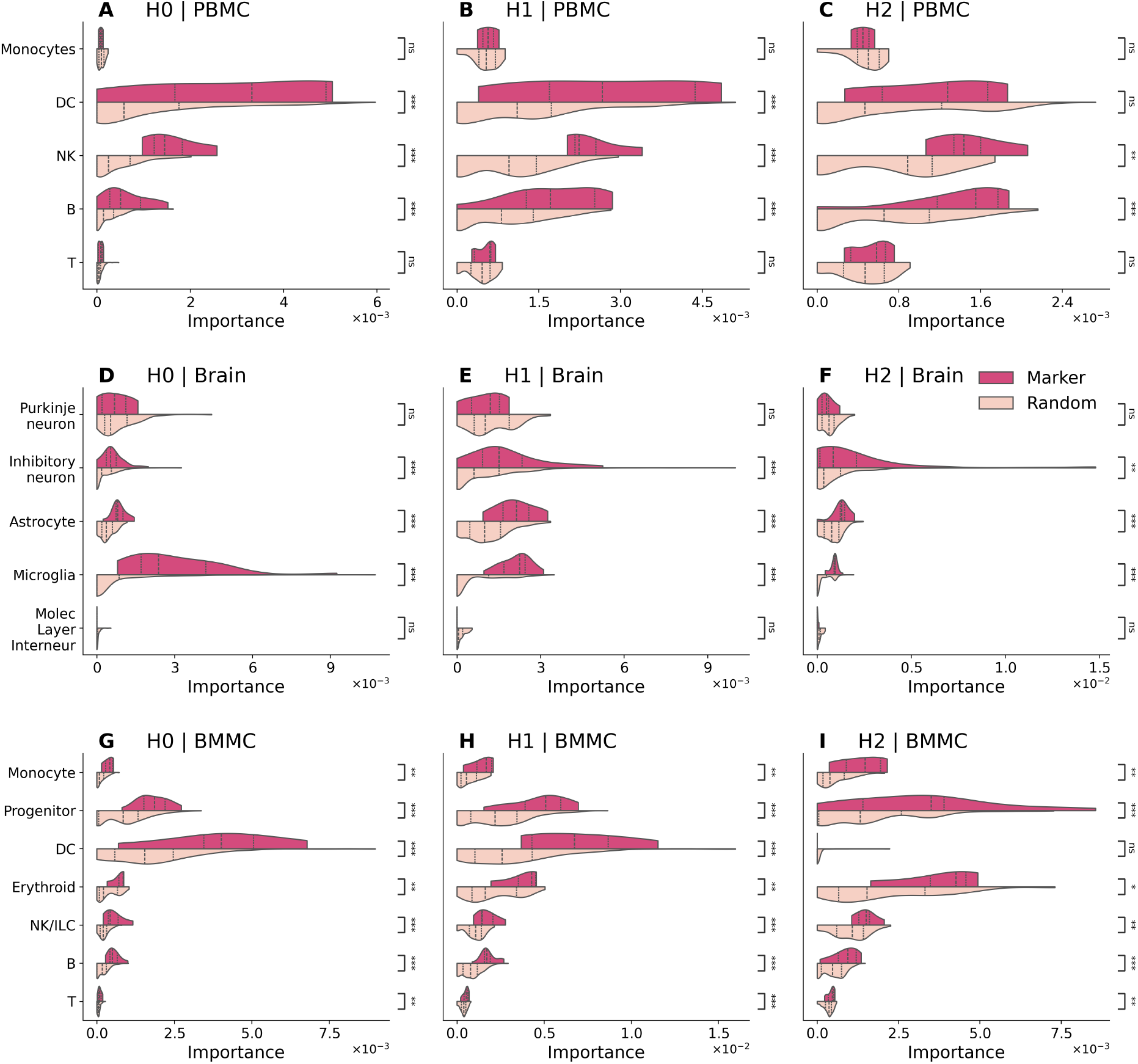
Biological marker genes have higher topological importance. Comparison of topology-importance scores between known marker genes and size-matched random gene sets across the three gene-expression data sets and homology dimensions *H*_0_, *H*_1_, and *H*_2_. Larger scores indicate a stronger effect of feature perturbation on the persistence diagram.

To assess the biological relevance of the features identified by XplainTDA, we interpreted the ATAC and GEX explanations independently. For ATAC, we obtained the peaks with the largest importance, and we linked these loci to transcription factors (TFs) by running motif analysis on the top-ranked peaks using HOMER [41] (Methods). For each considered cell type, we successfully identified biologically relevant TFs within the most important peaks (Fig. 5). In parallel, we performed gene set enrichment analysis on the top-ranked GEX features using the MSigDB database [42, 43] (see Additional file 2 and 3 for the complete list of motifs and GSEA results, respectively). Across the selected examples, we recovered FDR-significant gene sets consistent with the corresponding cell identities (Fig. 5). Importantly, in several cases the enriched transcriptional programs also converged with the TF independently identified from the ATAC peaks, providing complementary evidence from the two modalities that both the XplainTDA identifies valid biological features.

**Fig. 5:**
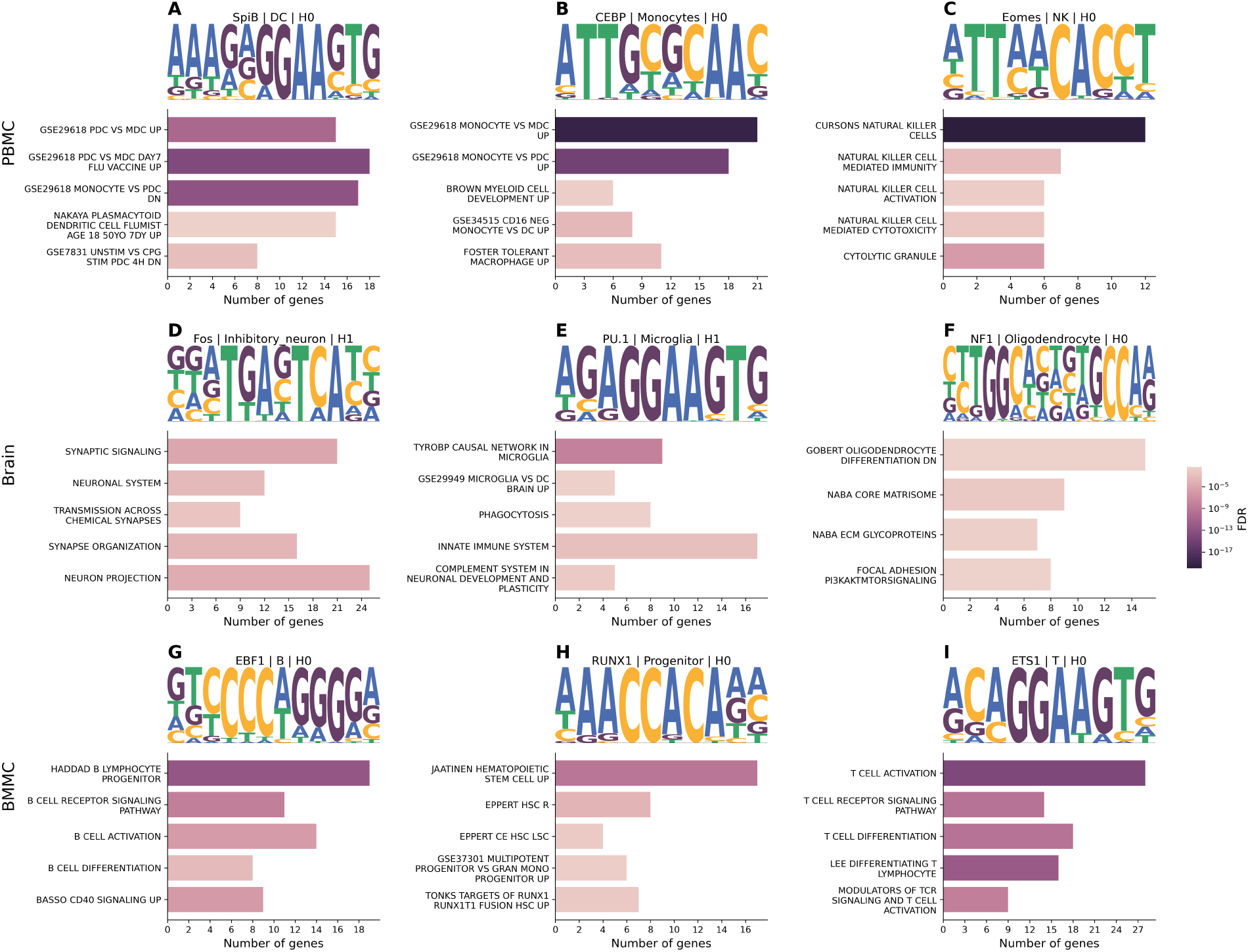
Cross-modal biological interpretation of selected XplainTDA explanations. Sequence logos show selected TF motifs identified by HOMER among the most important ATAC peaks for each cell type. Horizontal bar plots show selected FDR-significant gene sets from GSEA analysis in GEX genes for the corresponding cell type and homology dimension. Bar length indicates the number of overlapping genes and color represents the FDR on a logarithmic scale. The list whose names are structured as *GSE*/Family_name celltype_1 vs celltype_2 up/down* contain list that are up/down regulated in cell type 1 with respect to cell type 2 based on the finding of the research article indicate either with its GEO identifier or its first name of the publication.

For instance, in the PBMC data set, we identified motifs for Spi-B, C/EBP-family factors, and Eomes in DCs, monocytes, and NK cells, respectively (Fig. 5A–C). Spi-B is a key regulator of human plasmacytoid dendritic cells (pDC) development [44], C/EBP*α* promotes myeloid and monocyte lineage commitment [45], and Eomes has established roles in NK-cell development, maturation, and function [46]. The GEX enrichments showed biologically coherent agreement with these motifs. In DCs, several of the strongest selected gene sets specifically distinguished pDCs from myeloid DCs or monocytes, consistent with the requirement of Spi-B for pDC development [44]. In monocytes, we observed monocyte-specific signatures and a myeloid-development program derived from an experimental model of myeloid differentiation [47]. Similarly, NK cells were enriched for NK-cell identity and effector programs, including NK-cell-mediated cytotoxicity and the cytolytic-granule compartment. This agrees with the EOMES motif detected in the ATAC peaks, as Eomes is required for mature NK-cell homeostasis and cytotoxic function [48].

Among the motifs identified in the human brain data set, we found FOS, PU.1, and an NFI-family motif in inhibitory neurons, microglia, and oligodendrocytes, respectively (Fig. 5D–F). FOS is associated with activity-dependent neuronal transcription [49], PU.1 regulates human microglial gene expression and function [50], and NFI-family factors regulate oligodendrocyte-lineage differentiation and myelin-gene expression [51]. Consistently, inhibitory neurons were enriched for synaptic and neuronal programs, whereas microglia showed TYROBP-, phagocytosis-, innate-immune-, and complement-related enrichments. Oligodendrocytes showed oligodendrocyte-differentiation, extracellular-matrix, and focal-adhesion/PI3K–AKT–mTOR programs, consistent with established regulators of oligodendrocyte development and myelination [52, 53].

Finally, in the BMMC data set, we identified the EBF1 motif among the B-cell peaks. EBF1 is an essential regulator of B-cell lineage specification and modulates chromatin structure at B-cell-specific genes (Fig. 5G; [54]). Consistently, the GEX analysis recovered B-cell receptor signaling, B-cell activation, and B-cell differentiation programs. This provides a particularly direct correspondence between modalities, as EBF1 coordinates B-cell differentiation and is required for appropriate signaling in mature B cells [55]. The RUNX1 motif was found in hematopoietic progenitor peaks (Fig. 5H), consistent with its established role in the generation of hematopoietic stem cells [56]. Correspondingly, several of the selected GEX gene sets represented hematopoietic stem or progenitor programs. For instance, one enriched gene set comprised genes reported to be up-regulated following RUNX1–RUNX1T1 expression in normal human hematopoietic progenitors [57]. Among the T-cell peaks, we identified the ETS1 motif (Fig. 5I), while the GEX features were enriched for T-cell activation, T-cell receptor signaling, and T-cell differentiation programs, consistent with the established role of ETS1 in T-helper-cell differentiation and effector function [58].

## 4 Discussion

In this work, we introduced a perturbation-based framework to connect topological structure, derived from persistent homology, back to the molecular features that shape it (Fig. 1). Conventional feature-selection approaches, such as differential analysis or explainable AI, either look for statistical differences or interpret the outcome of a prediction model, respectively. Here, feature importance is instead defined by the extent to which a molecular perturbation alters the topology of the cell-state representation. This provides an additional perspective for studying biological organisation.

Across data sets and two molecular modalities, the inferred topology was reproducible over a broad range of *k*-nearest-neighbour graph sizes (Fig. 2 A – C) and with repeated cell subsampling (Fig. 2 D – F). These result prove the stability of PH under *k* – *NN* approximation and the its reproducibility within subsamples.

The biological relevance of the inferred topology was particularly evident for *H*_0_. This is not only the most stable homology dimension (Fig. 2) but, at an appropriate filtration scale, the connected components of the GEX topology closely reproduce the annotated cell-type structure (Fig. 3 D–F). This provides a direct link between the zero-dimensional topology of the learned manifold and established biological organisation. Persistent homology further extends this description beyond connectivity, with *H*_1_ and *H*_2_ capturing higher-order structures that are not directly represented by conventional clustering.

The main novelty of this paper is the perturbation-based explanation of the persistent structures. By masking individual genes or peaks and measuring the resulting change in the persistence diagram, XplainTDA assigns each molecular feature a topological importance score. Known marker genes consistently showed higher importance than random gene sets across all three data sets and homology dimensions *H*_0_, *H*_1_, and *H*_2_ (Fig. 4). Similarly, motif and GSEA analysis of most important peaks and genes recovered transcription factors with established roles in the corresponding cell populations and meaningful cellular pathways (Fig. 5). Together, these results indicate that changes in persistent topology can be traced back to biologically relevant molecular features.

Overall, our results support persistent homology as a practical tool for analysing the structure of single-cell data and show that its output can be made interpretable through in silico perturbation. XplainTDA combines these two ideas into a user-friendly Python package that identifies not only which persistent structures are present in a cell-state manifold, but also which genes or regulatory regions contribute most strongly to maintaining them.

## 5 Methods

### 5.1 Persistent homology and topological feature attribution

Let *X* ∈ ℝ^*N×P*^ denote the single-cell expression matrix, where *N* is the number of cells and *P* the number of genes. The data are first mapped to a low-dimensional representation

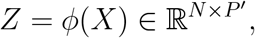

where *ϕ* denotes the fitted embedding model and *P* ^*′*^ *<< P*. When using the geometry-regularised autoencoder (GRAE) [9], an AE whose latent representation is constrained to preserve the geometry identified by PHATE [14] while retaining the out-of-sample mapping of an autoencoder. Given the expression matrix *X* ∈ ℝ^*N×P*^, PHATE first provides a low-dimensional reference representation

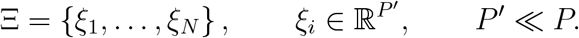

The autoencoder is then trained by jointly minimising the reconstruction error and the discrepancy between its latent representation *Z* and the PHATE embedding

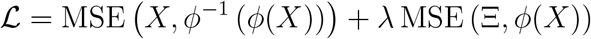

or, equivalently,

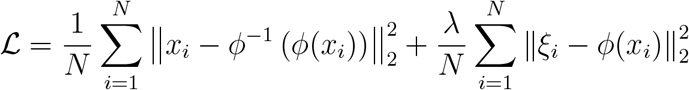

The hyperparameter *λ >* 0 controls the strength of the geometric regularisation: larger values constrain the latent representation more strongly toward the PHATE geometry. Persistent homology is computed from a weighted *k*-nearest-neighbor graph constructed in the embedding space. For each cell *z*_*i*_, the *k* nearest neighbours are identified using Euclidean distance, and the resulting directed neighbour relations are symmetrised to obtain an undirected graph *G* = (*V, E*) with edge weights

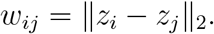

For a given distance threshold *ϵ*, we define the subgraph

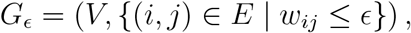

which contains only edges whose weight does not exceed *ϵ*. As *ϵ* increases, progressively more edges are added to the graph.

Each graph *G*_*ϵ*_ is expanded into its flag complex *K*_*ϵ*_, in which a set of vertices forms a simplex whenever all corresponding pairs of vertices are connected in *G*_*ϵ*_. Since increasing *ϵ* can only add edges and simplices

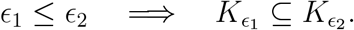

The resulting nested sequence of simplicial complexes is called a filtration, and persistent homology tracks the evolution of its topological features as *ϵ* increases [22, 23]. For each homological dimension *h*, the topology of *K*_*ϵ*_ is described by the homology group *H*_*h*_(*K*_*ϵ*_) [59]. Intuitively, *H*_*h*_ captures *h*-dimensional topological structures that are not filled in by higher-dimensional simplices. In particular, *H*_0_ describes connected components, *H*_1_ describes loops or tunnels, and *H*_2_ describes enclosed voids.

As *ϵ* increases, topological features can appear and subsequently disappear. Persistent homology tracks these changes across the filtration. A feature *l* is assigned a birth value *b*_*l*_, corresponding to the filtration value at which it first appears, and a death value *d*_*l*_, corresponding to the value at which it disappears by being filled by a higher-dimensional feature. Its persistence,

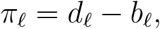

measures the range of filtration values over which the feature is present. Features with larger persistence, therefore, represent topological structures that are stable over a wider range of scales.

For each homological dimension *h*, the collection of birth and death values defines the persistence diagram

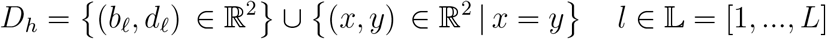

For computational cost, the set of *ϵ* values is defined by the list of distances in the *k* − *NN* adjacency matrix [34]; in this way, the algorithm applies only the steps that will definitely create or destroy a feature. Consequently, the last value for *ϵ* is simply the distance between the furthest apart neighbours. This implies that a topological feature may never experience death; the maximum filtration value replaces infinite death times. To quantify the contribution of individual biological features to the observed topology, each gene or peak *p* is perturbed across all cells, producing a modified expression matrix *X*^(*p*)^. The perturbation sets the expression of the selected feature to zero across cells. The embedding model is kept fixed throughout this procedure and is not retrained. The perturbed data are therefore projected using the fitted embedding model

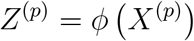

where *ϕ* is the trained GRAE model. Persistent homology is then recomputed from *Z*^(*p*)^ using the same neighbourhood and filtration parameters as for the original data, yielding the perturbed persistence diagram 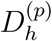.

The topological effect of gene perturbation is quantified using the q-bottleneck loss. For two persistence points *x* = (*b*_*x*_, *d*_*x*_) and *y* = (*b*_*y*_, *d*_*y*_), their matching cost is

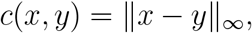

whereas the cost of matching a persistence point *x* to the diagonal is

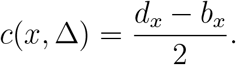

For a matching threshold *ϵ*, let *C*_*ϵ*_ denote the maximum number of off-diagonal persistence points that can be covered by an optimal matching whose cost does not exceed *ϵ*. The *q*-quantile bottleneck loss for two PD *D* and *E* is defined as

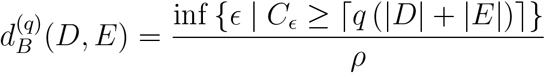

where *q* is the quantile of the distribution, and *ρ* is the diameter of the embedding, defined as

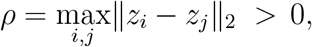

where *D* and *E* are the PD that include the diagonal, limited to the points that are matched to that.

We use *q* = 0.95, such that the loss corresponds to the smallest matching threshold required to account for at least 95% of the persistence points. This reduces the sensitivity of the score to a small number of unstable persistence features, enabling more robust and reliable estimation of gene and peak importance.

The importance of a gene or peaks *p* in homological dimension *h* is finally defined as

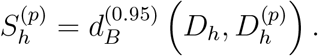

Genes are ranked independently within each homological dimension according to decreasing 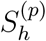. A higher score, therefore, indicates that perturbing a gene induces a larger change in the topology of the cell representation.

For the cell-type-specific explanation analyses, one GRAE was trained once per dataset and modality using an 85/15 training-validation split, then applied to all cells. The resulting embedding was then restricted to the cells belonging to each annotated cell type, and persistent homology and feature-perturbation scores were computed separately within each group while reusing the same fitted embedding model.

### 5.2 Stability with respect to the number of nearest neighbors

The stability of the PDs with respect to the number of neighbours used to construct the graph is assessed while the embedding is held fixed. A single embedding *Z* = *ϕ*(*X*) is first computed from the complete data set. Persistent homology is subsequently recomputed for a collection of neighbourhood sizes

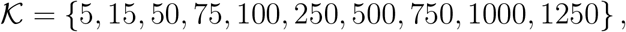

without refitting the embedding model. For each *k* ∈ *K*, a new weighted *k*-nearest-neighbor graph is constructed from the same *Z*, yielding persistence diagrams 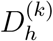 for each homological dimension *h*. Differences between persistence diagrams, therefore, reflect only changes in the neighbourhood graph, not in the learned representation. For zero-dimensional homology, the complete diagram 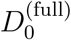 is computed using the complete Euclidean graph.

Stability across neighbourhood sizes is quantified by comparing persistence diagrams of all possible pairs of neighbourhood sizes, which are compared independently within each homological dimension,

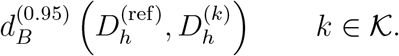

where `ref` is either the full-distance PD for *H*_0_ or *k* = 1250 for *H*_1_ and *H*_2_.

This analysis measures the extent to which the inferred persistent topology depends on the choice of *k*, while controlling for the embedding itself and using a common geometrical scale for comparisons.

### 5.3 Stability under cell subsampling

The stability of the persistent-homology representation with respect to the sampled cell population is evaluated by repeatedly recomputing persistent homology on stratified cell subsets based on cell-type annotations, while keeping the embedding model fixed. A single embedding of the complete data set *Z* = *ϕ*(*X*) is first computed and reused throughout the analysis.

For each replicate *r*, a fraction *α* of the cells is sampled independently within each group without replacement. If *N*_*c*_ denotes the number of cells belonging to group *c*, the number of cells sampled from that group is approximately 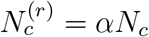, with at least one cell retained from each represented group. In the stability analysis, five stratified subsets are generated using *α* = 0.25. Let ℐ_*r*_ denote the indices of the cells selected for replicate *r*. Since the full embedding is kept fixed, the representation of each subset is obtained by restricting *Z* to the corresponding sampled cells

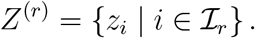

For each *Z*^(*r*)^, a new weighted *k*-nearest-neighbor graph is constructed and persistent homology is recomputed using the same filtration procedure as for the complete data set. This produces, for each replicate *r* and homological dimension *h*, a persistence diagram

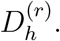

Stability is quantified by comparing every pair of independently sampled subsets. For replicates *r* and *s*, the *q*-bottleneck loss is computed separately for each homological dimension

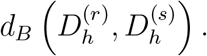

All pairwise comparisons are performed for every computed homological dimension. Because the same full-data embedding is used for all subsets, variation in these distances reflects the sensitivity of the persistent topology to cell sampling rather than stochastic variation in the dimensionality-reduction procedure. The visualisation of the persistence diagrams for the cell subsets is available in Fig. S1 (GEX) and Fig. S2 (ATAC).

### 5.4 Topological annotation from connected components

The connected components of the filtered *k*-nearest-neighbor graph provide a natural partition of the cells at each filtration value. Given the embedding *Z* = *ϕ*(*X*), the same weighted *k*-nearest-neighbor graph described above is constructed. For a filtration value *ϵ*, let

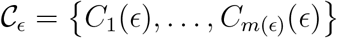

denote the partition of the cells into the *m*(*ϵ*) connected components of *G*_*ϵ*_. Equivalently, *C*_*ϵ*_ represents the partition associated with the zero-dimensional homology of the filtration at *ϵ*.

The graph edges are processed in increasing order of their filtration value. All edges with the same weight are introduced simultaneously, and the connected-component partition is retained only when the addition of these edges changes *C*_*ϵ*_. Thus, filtration values that do not alter the cell partition are discarded. This produces a sequence of informative filtration values

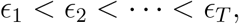

together with their corresponding component assignments.

To evaluate the agreement between the topology-induced partitions and an external cell annotation, let *Y* denote the reference cell-type labels and let *C*_*ϵ*_ denote the connected-component labels obtained at filtration value *ϵ*. For each retained filtration value, the agreement between *Y* and *C*_*ϵ*_ is quantified using adjusted mutual information (AMI); AMI corrects the mutual information for the agreement expected by chance

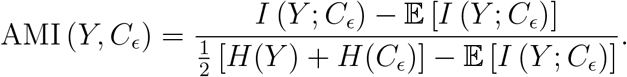

where *I*(·; ·) denotes mutual information and *H*(·) denotes Shannon entropy. AMI was computed using the *scikit-learn* implementation [60]. The filtration value providing the strongest correspondence with the external annotation is selected by maximising AMI

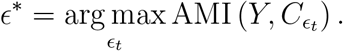

The connected components at *ϵ*^*∗*^ are then used as the topology-derived cluster assignment. This procedure, therefore, identifies the filtration scale at which the zero-dimensional topological structure of the embedding most closely matches the supplied biological annotation. We excluded the extremely small connected components, namely those with fewer than five cells, to reduce noise. Anyhow, the number of total cells excluded in each data set is always negligible. See Table S1 for the exact values of filtration, AMI, number of excluded cells and connected components.

### 5.5 Processing of the scRNA-seq and scATAC-seq data sets

The raw reads of the PBMC and Brain data sets were processed using *Cell Ranger ARC 2*.*0*.*2* [61], aligning them to the complete human genome (T2T) [62]. Their cell-type annotations and the pipeline to build the count matrices are taken from [19]. The gene expression count matrix of the BMMC was taken from [38], along with its cell type annotations. All matrices underwent the same preprocessing according to standard single-cell best practices [6]: quality control, filtering, library-size normalisation, and square-root transformation with *Scanpy* [63]. Doublets in the GEX data were identified with *Scrublet* [64]. The PBMC and Brain peak count matrices were built with *Episcanpy* [65] after *MACS2* [66] peak calling, also using the code from [19]. In both scRNA-seq and scATAC-seq matrices, we kept only the highly variable genes or peaks. For the code to reproduce the preprocessing, see https://github.com/gmalagol10/XplainTDA.git. We ran the GRAE independently for each count matrix, setting the latent space dimension as the cubic root of the input dimension, as suggested in [19]. Finally, we stored both GEX and ATAC count matrices in one unique *Muon* object [67]. For the detailed preprocessing from FASTQ files to count matrices see [19] for the PBMC and Brain data set, and [68] for the BMMC one.

### 5.6 GEX markers and ATAC motif analysis

The GEX markers for the PBMC and BMMC data sets were taken from [38], whereas for the Brain data we used [69–76]. The complete list of markers is available in Table S2. Since the count matrices contain only highly variable genes, not all suggested markers are present in the GEX list; therefore, we excluded test instances with fewer than one marker. To control for differences in gene abundance and detectability, null genes were matched to marker genes according to their global mean expression and detection coverage. Genes were assigned to rank-based expression/coverage bins, and matched control genes were sampled from the corresponding bins, excluding known marker genes. For each cell type and homology dimension, we compared the mean XplainTDA importance of the marker set with a null distribution generated from 10000 matched random gene sets of equal size, sampling one matched control per marker. One-sided empirical *p*-values were computed as the fraction of null sets with importance at least as large as the observed marker set and were corrected using the Benjamini–Hochberg FDR procedure [77]. Statistical annotations were displayed using *statannotations* [78]. The motif analysis was performed with HOMER [41]. For each cell type and homological dimension, we selected the top 200 ATAC peaks ranked by their topological importance score and performed motif enrichment using the T2T reference for the PBMC and Brain data sets, and hg38 for BMMC.

To functionally interpret the GEX explanations, we performed gene-set enrichment analysis on the top 100 genes ranked by their topological importance score for each cell type and homological dimension. We tested gene sets from the MSigDB v2026.1 Hallmark (H), C2, C5, and C7 collections [42, 43] using a hypergeometric test, followed by Benjamini–Hochberg correction for multiple testing [77]. For visualization, we selected up to five FDR-significant, non-redundant gene sets per representative cell type, prioritizing terms that were biologically consistent with the cell identity and, where applicable, with the TF identified independently by the ATAC motif analysis.

## 6 Availability of Data and Materials

Code and tutorial for XplainTDA are available at https://github.com/gmalagol10/XplainTDA.git. The code to reproduce all the results is available at https://github.com/gmalagol10/XplainTDA/tree/main/reproducibility.

## 7 Acknowledgments

We thank Dr Mafalda Dias and Dr Jonathan Frazer for their valuable inputs when starting the project. We thank Prof. Guy Wolf for helpful suggestions on validating persistent homology in single-cell data. We thank Prof. Christina Kuttler for serving as the university supervisor for I.G.’s master’s thesis, from which this work developed, and for meeting with us to discuss the project. We thank the BMC Bioinformatics Core Facility for providing access to their HPC cluster. Figure 1 was partially created with BioRender.

## 9 Contributions

G.M., K.T.S. and M.C.T. designed the study. I.G. and G.M. implemented the method and performed the analysis. I.G., G.M. and M.C.T. wrote the manuscript with help from K.T.S. All authors reviewed and approved the manuscript.

## 10 Declarations

The authors declare no conflict of interest.

## 11 Supplementary files

The full list of marker genes, their references, and supplementary tables are available in “Additional file 1”. The complete list of motifs identified with HOMER is shown in “Additional file 2”. The supplementary figures are available in the file “Additional file 3”.

## Additional file 1: Supplementary tables

**Table S1:** Metadata regarding the procedure to retrieve the cell type annotation from the connected components.

| Data set | Filtration value | AMI | Number of cells | Number of removed cells | Number of removed CC |
| --- | --- | --- | --- | --- | --- |
| PBMC | 0.10229071369 | 0.8819 | 5639 | 61 | 48 |
| Brain | 0.08106958423 | 0.8005 | 2568 | 104 | 70 |
| BMMC | 0.09515247711 | 0.7368 | 5577 | 243 | 200 |

**Table S2:**
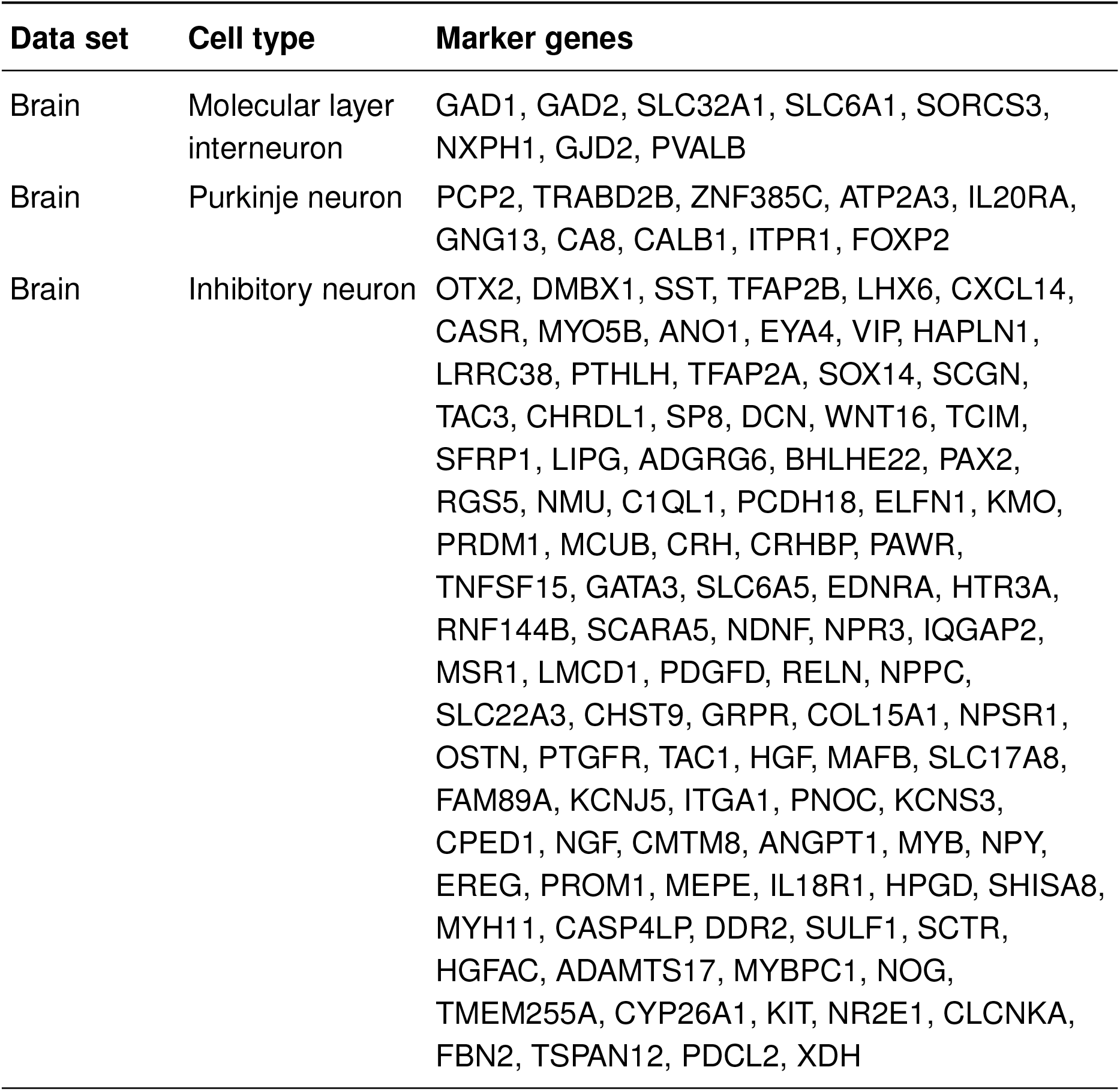
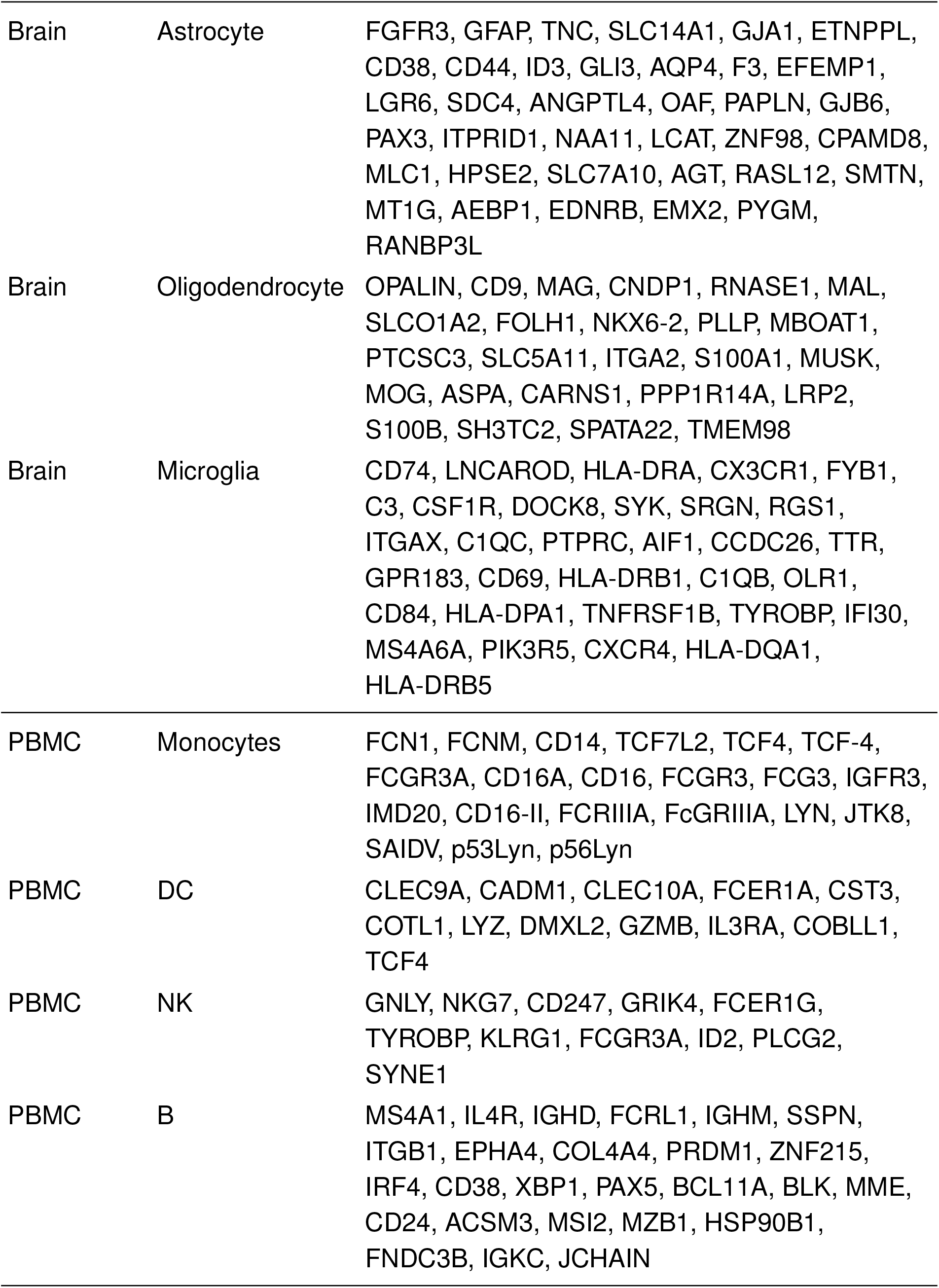
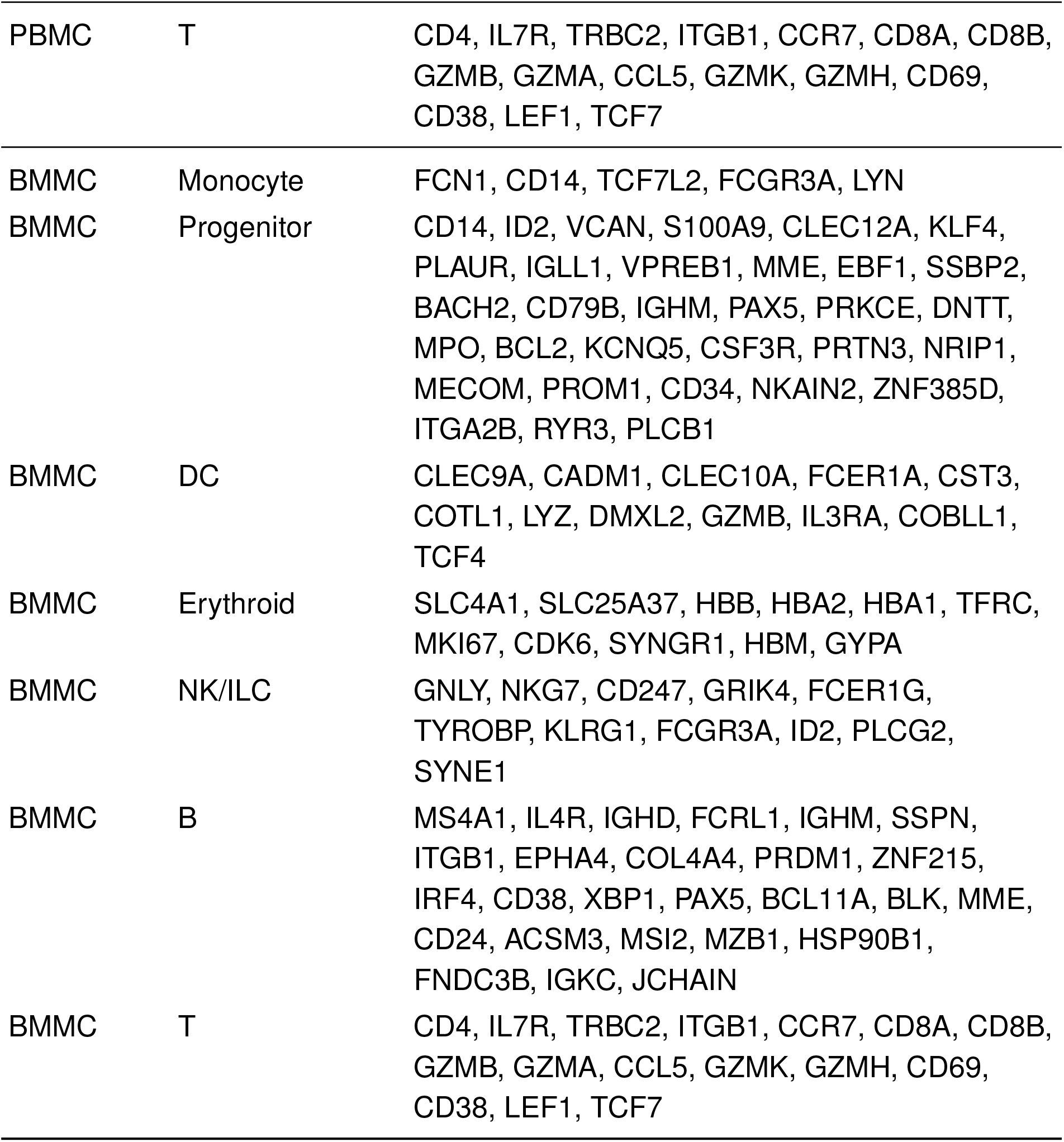
Marker genes used for the comparison between topologically important features and known cell-type markers across the three gene-expression data sets.

## Additional file 3: Supplementary figures

**Fig. S1:**
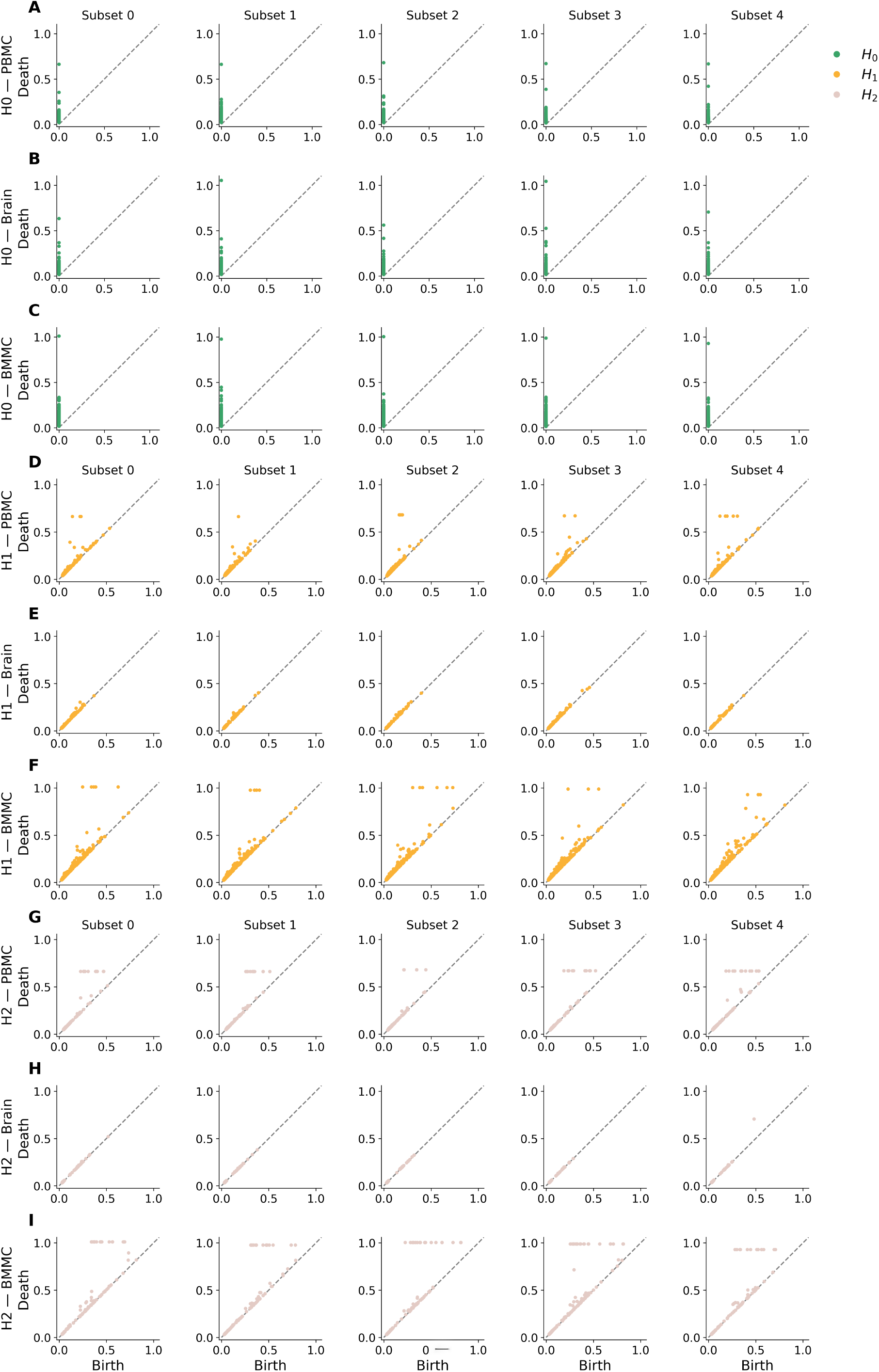
Persistence diagrams of all the subsets of cells in the GEX space.

**Fig. S2:**
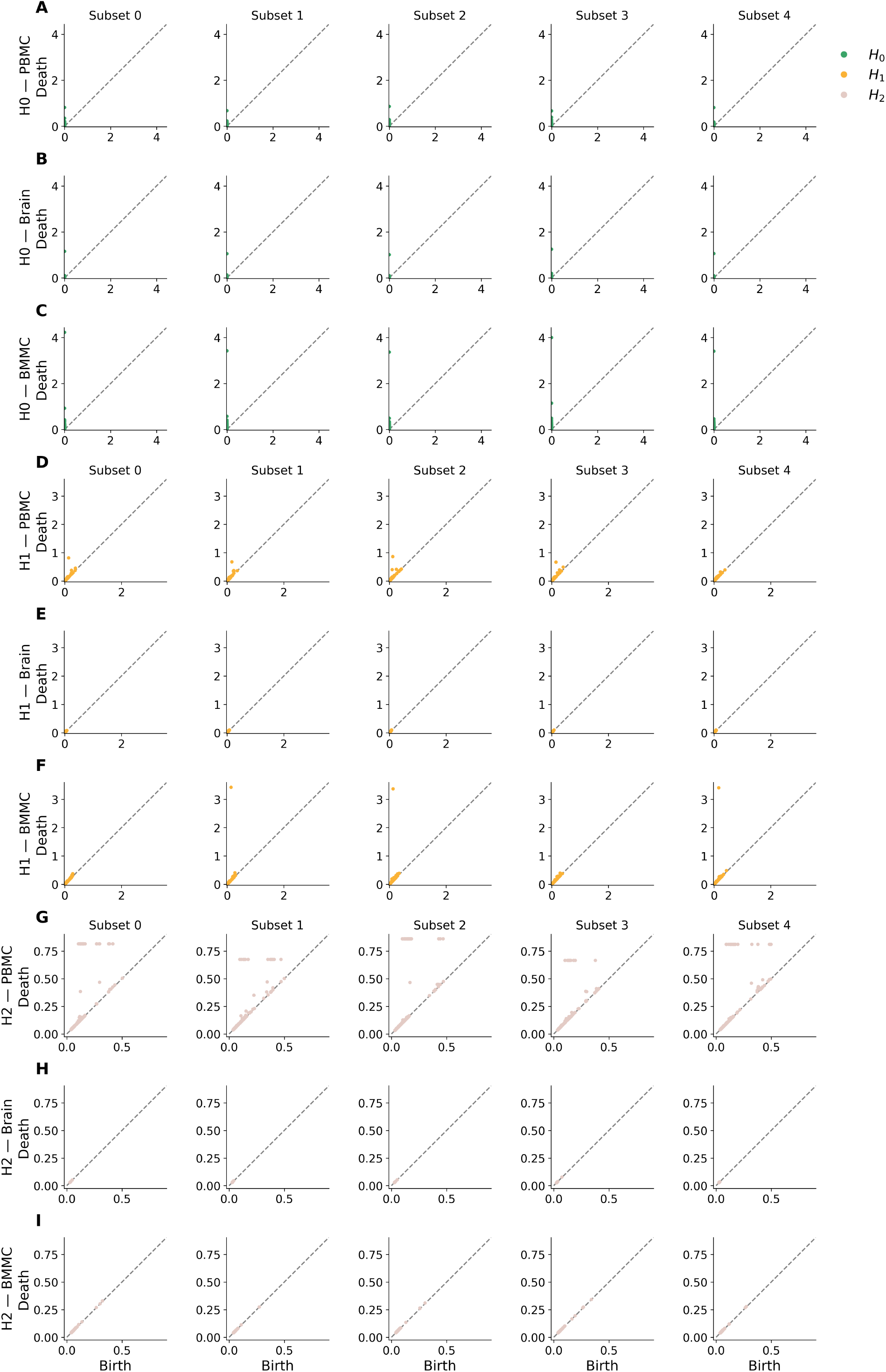
Persistence diagrams of all the subsets of cells in the ATAC space.

